# Hippocampal extracellular matrix remodeling and temporally distinct glial reorganization following peripheral injury

**DOI:** 10.64898/2026.09.30.755817

**Authors:** Chuang Ge, Vaneeza Kausar, Lele Xu, Amirabbas Maghsoudi, Gabriella Shtudland, Michael Amrami, Michael Betancourt, Areej Niaz, Nuzhat Anchal, Tiffany Grapsas, Jacob Zirkiyev, Sebastian Alvarado, Rinat Abzalimov, Ye He, Maral Tajerian

## Abstract

Chronic pain is frequently accompanied by anxiety and other affective disturbances, but the molecular and cellular changes linking peripheral injury to persistent alterations in brain function remain incompletely defined. The hippocampal extracellular matrix (ECM) regulates synaptic plasticity, cellular signaling, and neuroimmune interactions and may therefore contribute to pain-related central plasticity. We used a tibial fracture/cast immobilization model to examine temporal changes in hippocampal ECM, glial morphology, and affective behavior following peripheral injury. Behavioral testing was performed at 3, 7, and 20 weeks after injury, while high-dimensional morphometric analysis of microglia and astrocytes was performed at 1, 3, 7, and 20 weeks. Hippocampal ECM-associated molecular signals were mapped at the chronic stage using decellularization coupled with MALDI-TOF mass spectrometry imaging. Injured mice developed mechanical hypersensitivity and reduced exploration in two anxiety-related paradigms during the early and chronic stages, with these behavioral phenotypes no longer detected at 20 weeks. At 7 weeks, injury was associated with changes in hippocampal ECM-associated peptide signals, including reduced signals corresponding to four ECM-associated proteins. Microglia and astrocytes exhibited distinct, time-dependent changes in density and multidimensional morphology, with some alterations persisting after behavioral measures returned toward control levels. In vitro, BV2 microglia exhibited rigidity-dependent changes in morphology, phagocytic behavior, and calcium signaling. Together, these findings define hippocampal ECM–glial phenotype following peripheral injury and identify matrix biomechanics as a candidate regulator of microglial function.

## Introduction

Chronic pain is increasingly regarded as a disorder of maladaptive central plasticity rather than a persistent peripheral nociceptive condition[1, 2]. Neuroimaging and translational studies have established that chronic pain involves large-scale reorganization of limbic and affective circuits, including the amygdala, hippocampus, and prefrontal cortex[3–6]. Structural and functional alterations within these circuits correlate with affective symptoms, altered motivational states, and impaired cognitive flexibility[7], suggesting that chronic pain reflects sustained central plasticity rather than ongoing tissue pathology alone. The accompanying high prevalence of anxiety and depression in chronic pain patients suggests that mechanisms underlying pain chronification may engage brain regions relevant to affective regulation[8, 9].

Mouse models of bone injury have shown evolving nociceptive and physiological signs of pain, with injured mice displaying nociceptive sensitization, edema, and warmth, alongside altered peripheral neural signaling and cytokine generation[10, 11]. In addition, complex behavioral analyses have demonstrated signs of anxiety and memory impairment, paralleled by structural and biochemical changes in select brain regions such as the amygdala, perirhinal cortex, and hippocampus[12]. This raises the possibility that central immune mechanisms contribute to the maintenance of pain-related behavioral comorbidities.

The hippocampus plays a key role in integrating stress, memory, and emotional processing. Both astrocytes and microglia are increasingly recognized as active regulators of synaptic plasticity and neuroimmune signaling, rather than passive bystanders[13]. Astrocytes shape synapse formation, transmission, and memory-related network activity[14], whereas microglia, beyond their classical immune surveillance role, regulate synaptic refinement, trophic factor release, inflammatory signaling, and circuit homeostasis[15]. These glial functions are dysregulated in chronic pain[16], and, consistent with this, persistent inflammatory pain disrupts hippocampal synaptic plasticity and impairs spatial memory through microglia-dependent BDNF and proBDNF signaling[17].

Although inflammatory cascades in chronic pain have been extensively studied, comparatively less attention has been paid to the extracellular environment in which these processes unfold. The brain extracellular matrix (ECM) shapes synaptic stability, constrains diffusion, and influences neuronal–glial interactions[18–20]. ECM components such as laminins, tenascins, and perineuronal net constituents undergo activity-dependent remodeling and can modulate synaptic efficacy and circuit stability. Such structural substrates may therefore provide a regulatory framework for neuroimmune plasticity during pain chronification[21, 22].

Glia and the ECM shape one another. Astrocytes supply matrix components and microglia remodel them through proteolytic and inflammatory pathways[23], while matrix stiffness and composition in turn set glial activation state through mechanosensitive signaling[24, 25]. The consequences for neurons are best documented in the hippocampus, where excitotoxic injury triggers tissue plasminogen activator/plasmin-mediated degradation of laminin. Laminin loss precedes neuronal death, and preserving the neuron–laminin interaction protects hippocampal neurons[26]. The same proteolysis leaves the induction of long-term potentiation intact but abolishes its maintenance[27], implicating laminin–cell contact in sustained rather than initiated plasticity. Glycoprotein remodeling constrains glia in a comparable way: fibronectin aggregates persisting in chronically demyelinated multiple sclerosis lesions block oligodendrocyte progenitor differentiation and remyelination[28], and the laminin-, tenascin-, and proteoglycan-rich matrix deposited by reactive astrocytes after focal CNS injury limits axonal regrowth[29].

Moreover, matrix-dependent signaling pathways influence microglial activation states. Matrix-associated inflammatory modulation has been observed in microglial responses to LPS [30], and recent studies indicate that matrix composition can modulate microglial reactivity and mechanosensitive pathways [31]. How the molecular composition of the hippocampal ECM is altered at the chronic pain stage, and how such changes relate to the evolving glial response, remains poorly defined. We previously reported that manipulation of the hippocampal ECM alters pain and pain-related memory in this same tibial fracture model [29]; however, the temporal evolution of ECM-glia reorganization in this system has not been characterized.

We previously showed, in this same tibial fracture model, that the chronic pain stage involves remodeling of the hippocampal ECM in the form of reduced matrix rigidity, decreased aggrecan, and increased MMP8; furthermore, downregulating the injury-induced MMP8 was sufficient to relieve tactile hypersensitivity and pain-related memory deficits, establishing the hippocampal ECM as a causal regulator of the chronic pain phenotype[32]. Here we extend that work by mapping the molecular composition of the injured hippocampal ECM at the chronic stage with MALDI-TOF imaging, tracking microglial and astrocytic morphology across multiple post-injury time points, and testing in vitro whether substrate rigidity shapes microglial state.

Here, we test the hypothesis that chronic peripheral injury remodels the molecular composition of the hippocampal extracellular matrix at the chronic pain stage, in parallel with temporally dynamic changes in glial morphology and affective behavior. We combined longitudinal behavioral assays and high-dimensional morphometric analysis with mass spectrometry–based molecular mapping at the chronic pain stage to examine how structural matrix alterations relate to neuroimmune plasticity in this model. We further used an *in vitro* biomechanical platform to examine whether substrate rigidity is associated with changes in microglial morphology, phagocytic behavior, and calcium signaling, and how these rigidity-dependent responses are modulated by LPS stimulation.

## Methods

### Animals

Male C57BL/6J mice ages 12–14 weeks were purchased from a commercial supplier (Jackson Labs, Sacramento, CA, USA) and were allowed to habituate to the animal facility for a minimum of 10 days prior to the experiments. Mice were housed in groups of 2-4 on a 12-hr light/dark cycle and an ambient temperature of 22 ± 3°C, with food and water available *ad libitum*. All animal procedures and experimental designs were approved by the *Queens College Institutional Animal Care and Use Committee* (Flushing, NY, USA) and followed the “animal subjects” guidelines of the *International Association for the Study of Pain*.

### Tibia Fracture and Cast Immobilization

Mice were randomly allocated to either the control or the fracture/cast group. Animals were anesthetized with 1.5% isoflurane and a hemostat was used to induce a closed fracture of the right tibia just distal to the middle of the tibia and the hindlimb was wrapped in casting tape (cat. # 82001, Scotchcast^TM^ Plus, 3M, UK), as previously described[33]. After the procedure, the mice were given subcutaneous buprenorphine (0.05 milligrams per kilogram [mg/kg]) and enrofloxacin (5 mg/kg) for the next two days, as well as normal saline (1.5 ml once) for post-operative analgesia, prevention of infection and prevention of dehydration. Mice were inspected daily to ensure the cast was positioned properly through the 3-week period of cast immobilization. Mice were provided with chow pellets postoperatively *ad libitum*; dietary gels were also made available on the cage floor for mice having undergone surgery. Casts were removed 3 weeks after surgery under brief isoflurane anesthesia.

Behavioral assessments were performed at 3, 7, and 20 weeks post-surgery (early, chronic, and recovery stages, respectively). Brains for immunohistochemical and morphological analysis were collected at 1, 3, 7, and 20 weeks post-surgery. Brains for MALDI-TOF imaging were collected at 7 weeks post-surgery (chronic stage).

### Behavioral testing

All analysis was blinded to the identity and experimental condition of the animal. Mice were habituated to handling the experimenter for a few minutes each day for 7 days before initiation of the behavioral tests.

#### Mechanical sensitivity

Calibrated monofilaments (Stoelting Co., USA) were applied to the plantar surface of the hind paw and the 50% threshold to withdraw (grams) was calculated as previously described[34]. The stimulus intensity ranged from 0.004 to 1.7g, corresponding to filament numbers 1.65, 2.36, 2.44, 2.83, 3.22, 3.61, 3.84, 4.08, 4.17, and 4.31. For each animal, the actual filaments used within the aforementioned series were determined based on the lowest filament to evoke a positive response (response = flexion reflex) followed by five consecutive stimulations using the up–down method. The filament range and average interval were then incorporated along with the response pattern into each individual threshold calculation.

#### Y-maze

Rodents’ natural inclination to explore new environments was evaluated by quantifiying spontaneous alteration in exploring the arms of the maze[35]. In a Y-shaped maze with three identical arms (A, B, and C), mice tend to choose a new arm over one they’ve already visited. The maze was made in-house using opaque white acrylic. Mice were placed in the center and allowed to explore for 5 minutes. Mice with strong spatial working memory typically entered a new arm without revisiting a previous one. The test measured spontaneous alternation by tracking unique sequences of arm entries (e.g., ABC, BCA). The percentage of alternation was calculated by dividing the number of unique sequences by the total arm entries minus two.

#### Zero maze

The apparatus was made in-house and had the following dimensions: inner circle diameter=47 cm, outer circle diameter=54 cm, height =25cm. Mice were placed at the boundary between the open and enclosed regions and allowed to explore freely for six minutes. The time spent by the mice in each of these regions was quantified[36].

#### Open field

The apparatus was built in-house as a 28X28X28 (LXWXH) acrylic cube. Mice were placed in the center of the field and allowed to explore for 6 minutes. The time spent in the central 25% area vs the entire arena was calculated in addition to the speed of locomotion in the central and peripheral areas[37].

### MALDI-TOF

The Matrix-Assisted Laser Desorption/Ionization Time of Flight (MALDI-TOF) Mass Spectrometry Imaging (MSI) was carried out in the MALDI MS Imaging Joint Facility at the Advanced Science Research Center of City University of New York. High-purity grade trypsin, alpha-cyano-4-hydroxycinnamic acid (CHCA), acetic acid, acetonitrile, and Phosphorus (red) were purchased from Millipore Sigma-Aldrich (USA). Optima UHPLC/MS-grade acetonitrile, trifluoroacetic acid (TFA), ammonium bicarbonate, Sodium Dodecyl Sulfate (SDS), and water were purchased from Fisher Scientific (USA). n = 1 animal per condition; left and right hippocampi were treated as spatial/technical measurements within the same biological sample.

#### Tissue sectioning

Mouse brains were harvested and snap-frozen for 5 min on an aluminum boat floating on liquid nitrogen. The frozen tissue was cryosectioned at 12 μm thickness using Minux® FS800 Cryostats (RWD, China) at −18 °C set for specimen head and −20 °C for the chamber. The tissue cryosections were gently transferred onto the pre-cooled conductive side of indium tin oxide (ITO)-coated glass slides (Bruker Daltonics, Bremen, Germany) and thaw mounted using finger. Sample slides were stored at −80 °C until further usage. For each mouse, spectra from both the left and right hippocampi were collected and included in the quantitative analysis.

#### Decellularization and washing

Tissues were decellularized by incubating in a 0.01% SDS at 4 °C for 48h, with slight shaking. Then tissues were rinsed three times in a 10 mM ammonium bicarbonate solution at 4 °C for 15 minutes to remove excess SDS. A six-step rinse protocol for tissues: 70% ethanol, 100% ethanol, Carnoy’s fluid, 100% ethanol, water, and 100% ethanol. Carnoy’s fluid was prepared by mixing 60 ml of ethanol, 30 ml of chloroform, and 10 ml of acetic acid. Each rinse step lasted 30 seconds, except for Carnoy’s fluid step, which lasted 2 minutes[38].

#### Trypsin digestion

Mounted cryosections on ITO slides were desiccated in vacuum for 45 min at room temperature followed by trypsin deposition using Hamilton syringe pump (New Era Pump Systems, Inc). A trypsin solution of 0.1 µg/µl in 50 mM ammonium bicarbonate buffer/acetonitrile (90/10, v/v) was deposited onto brain sections through HXT M5 sprayer (HTX LLC., USA) at a flow rate of 8.0 µl/min, a nozzle temperature of 30 °C, and a heated tray temperature of 30 °C for 12 cycles without drying time between the cycles. A spray velocity of 750 mm/min, track spacing of 2 mm, N_2_ gas pressure of 10 psi and flow rate of 3 L/min, and nozzle height of 40 mm were used. A humidity chamber was created inside petri dishes using a KimWipe sprayed with 2 mL of the 50 mM ammonium bicarbonate buffer/acetonitrile (90/10, v/v) solution. Following trypsin application, the slides were incubated in the humidity chamber at 37 °C for 18 hrs.

#### Matrix Deposition

Tryptic digestion was terminated as the slides were removed from the hydration chamber and deposited with matrix using HTX M5 sprayer (HTX LLC., USA). Matrix CHCA prepared in a solution of 5mg/ml in acetonitrile/TFA/water (90/0.1/9.9, v/v/v) was deposited at a flow rate of 0.05 ml/min and a nozzle temperature of 85 °C for 6 cycles without drying time between the cycles. A spray velocity of 700 mm/min, track spacing of 2 mm, N_2_ gas pressure of 10 psi and flow rate of 3 L/min and nozzle height of 40 mm were used.

#### Spectral processing and statistics

The MSI data from FlexImaging were converted and uploaded into the SCiLS Lab software (SCiLS–Bruker Daltonik GmbH, Germany) used for pipeline preprocessing and statistical analysis. The preprocessing pipeline applied baseline removal using the iterative convolution algorithm with sigma = 20 and iterations = 20. Subsequently, the total ion current (TIC) of every spectrum was normalized to one. Principal component analysis (PCA) of MALDI-TOF spectra was performed in SCiLS to evaluate the multidimensional distributions of the identified peaks from injured and control hippocampal regions. Spectra from the injured hippocampus and control hippocampus were processed individually, with normalization using unit variance scaling and minimal interval binning of ±0.200 Da. PCA was conducted with 3 principal components to capture the primary variance in the data. Peaks from the injured and control regions were visualized as red and black dots, respectively, to highlight differences in their spectral profiles.

The intensity of each scan over the entire acquired mass range was mapped on the left and right hippocampus region within each brain section to visualize the location of each m/z value detected. These images were generated and visualized using SCiLS. Box plot summaries displayed median intensity, interquartile range, and 0–99% whiskers, with outliers identified as spots whose intensities fell outside this range.

#### Compounds Identification/Annotation

The spectra were manually interpreted. Initially, the elemental compositions of each detected ion were predicted using Metaboscape (Bruker Daltonics, Germany), based on precise mass measurements (with a mass accuracy of less than 5 ppm) and corresponding isotopic patterns. These ions were subsequently cross-referenced with several metabolite databases, including The Human Metabolome Database (HMDB), METLIN, LIPID MAPS, as well as a custom-built metabolite library derived from previous LC-MS experiments. The obtained metabolite identification was further cross-referenced with findings from previously published studies based on both m/z value and spatial distribution in rodent brain[39]. ECM analysis was performed using peptideMass (https://web.expasy.org/peptide_mass/) to predict the masses of peptides generated from extracellular matrix (ECM) proteins after trypsin digestion. The generated peptide masses were compared to m/z values obtained from MALDI-TOF MS, with confirmation of ECM proteins when three or more peptides matched MALDI results and had similar expression patterns.

### Immunohistochemical staining and analysis

Mice were anesthetized by 300 μl of ketamine/xylazine cocktail, followed by transcardiac perfusion with 0.9% saline solution. Brains were carefully removed, postfixed in 4% paraformaldehyde for 2 days, and rinsed in phosphate-buffered saline for 3 days. Contralateral brain hemispheres were then embedded in agarose, and cross-sections were cut at room temperature at 40-μm thickness on a vibratome (Leica VT 1200S; Leica Biosystems, USA). Ten sections per mouse were randomly chosen for staining. Free-floating immunohistochemistry was done by using tris-buffered saline + 0.1% Tween-20 (TBST) as the wash buffer and 10% donkey normal serum (cat. #ab7475, Abcam, USA) in phosphate-buffered saline as the blocking buffer. A permeabilization step was added (before blocking and staining) using 0.1% triton and 0.6% hydrogen peroxide in 1X TBST. The following primary antibodies were used (diluted in blocking buffer): mouse monoclonal anti-glial fibrillary acidic protein (GFAP; 1:5000, cat. MAB3402, Millipore, USA) and rabbit polyclonal anti-ionized calcium-binding adapter molecule 1 (Iba1, 1:800, cat. 019-19741, Wako, Japan). The following secondary antibodies were used (diluted in 1X TBST): donkey anti-mouse immunoglobulin G (IgG, H + L), Alexa Fluor 647 (AF647, 1:500, cat. ab150107, Abcam), and donkey anti-rabbit IgG (H + L) Alexa Fluor 488 (AF488, 1:500, cat. 711-545-152, Jackson Immunoresearch, USA). Brain sections were mounted on slides using fluoromount aqueous mounting medium (cat. F4680, Sigma,). Images were analyzed as a 3-μm step z-stack of 10 slices (20X objective magnification) using fluorescent imaging (Keyence BZ-X700, USA). Two sections per mouse were randomly chosen for analysis. GFAP+ and Iba1+ cells were manually quantified in each z-stack using the “Mark and Count” tool, and cell area were measured in image J (National Institute of Health, USA).

#### Single-Cell Morphological Quantification

Single-cell morphological analysis was performed using a semi-automated ImageJ-based workflow adapted from Kim et al. (2024). Briefly, fluorescent images were converted to 8-bit format, binarized, and thresholded using dataset-specific parameters optimized to preserve full cellular morphology while minimizing background noise. Individual cells were segmented based on predefined area criteria to exclude artifacts and overlapping structures. Skeletonization was performed using the AnalyzeSkeleton plugin in ImageJ, and additional morphometric parameters were extracted using the FracLac plugin. A total of 27 morphological features were quantified per cell, including measures of branching complexity (e.g., number of branches, junctions, and endpoints), branch length, territory span, area, circularity, and density. The same analytical pipeline was applied to GFAP-stained astrocytes with optimized thresholding and area-filtering parameters to account for differences in astrocytic morphology.

#### Data Preprocessing and Dimensionality Reduction

Morphological feature tables were exported to Excel and analyzed in R (version 4.3.1). Non-feature metadata columns (ROI_ID, MouseID, Model, Date) were excluded before analysis, and all morphological features were standardized (z-score normalization). Principal component analysis (PCA) was performed using the prcomp function, retaining the smallest number of principal components that together accounted for ≥ 90% of the total variance.

#### UMAP Visualization and Unsupervised Clustering

Cells were partitioned into four morphological states by k-means clustering (k = 4, nstart = 50; set.seed(123)) applied to the retained principal components. The number of clusters was fixed a priori; silhouette analysis across k = 2–10 identified no discrete optimum in either cell type, indicating that glial morphology varies along a continuum rather than forming separable subtypes, and k = 4 was therefore used as a partition of this continuum. Because k-means labels are arbitrary, clusters were renumbered by descending size to ensure reproducible indexing. UMAP (uwot package; n_neighbors = 15, min_dist = 0.1, Euclidean distance; set.seed(123)) was applied to the same principal components for two-dimensional visualization only; UMAP coordinates were not used for clustering or for any statistical comparison. The identical preprocessing, dimensionality-reduction, and clustering pipeline was applied separately to Iba1⁺ microglia and GFAP⁺ astrocytes.

### Cell Culture

BV-2 cells derived from murine neonatal microglia were used as a substitute for primary microglia. All cells were plated in 6-well collagen-coated plates except for the controls. The 6-well plates were primed according to the manufacturer’s instructions (Cat # 5190, Advanced Biomatrix) by using bovine collagen Type 1 solution (Cat # 5005, Advanced Biomatrix) and were incubated for 1 h prior to the experiments. After the incubation period, the plates were washed twice with 1X Dulbecco’s Phosphate Buffered Saline (DPBS), followed by adding growth medium: Dulbecco’s Modified Eagle Medium, supplemented with 10% FBS, 1% penicillin & streptomycin to avoid dehydration (DMEM +/+). Cells were seeded at 2.5 × 10⁵ cells per well in 6-well collagen-coated plates with 3 mL of culture medium and were allowed to adhere and grow at 37°C, 95% humidity, and 5% CO₂ for 24 hours. The culture medium was then changed, and LPS (Cat # 00-4976-93, ThermoFisher) was added to the respective plates at a final concentration of 2 µg/mL.

#### Cell death assay

After 24 h of incubation with LPS, the supernatant from each well was collected and labeled. The wells were then washed with 1×DPBS. To detach adherent cells, 2 mL of 0.25% trypsin was added to each well, and the plates were then incubated for 7 minutes. Then, 2 mL of DMEM +/+ was added to neutralize trypsin, and the cell suspension was collected. The collections were added to their respective collection tube and centrifuged for 7 minutes at 1500 RPM. The collection tubes were decanted, and the pellets were resuspended in 200 µl of DMEM +/+. 20 µl of each sample tube was mixed with 20µl of 4% Trypan Blue and kept on ice for 5 minutes before analysis. The live and dead cells were distinguished and counted using a hemocytometer and an Olympus CX31microscope.

#### Morphology assay

Following 24 hours of incubation, non-adherent cells were removed by washing with 1×DPBS. Adherent cells were fixed with 4% formaldehyde (1.5 mL per well) for 11 minutes at 37° C with 95% humidity and 5% CO₂. After fixation, the cells were washed again with 1×DPBS and imaged on a Keyence BZ-X810 microscope using a 20×/0.75 objective lens. BV-2 cell morphology was quantified using ImageJ (Fiji) with a Cellpose plugin to obtain area and roundness measurements. The data were analyzed using GraphPad Prism.

#### Phagocytosis assay

After 24 h of incubation of Cytosoft plates, as described above, 2 µm carboxylate-modified microspheres (F8827, Thermofisher) were added to the wells at a concentration of 5 beads per cell. The cells were then incubated at 37° C with 95% humidity and 5% CO₂ for 120 minutes. After incubation, the wells were fixed using 1.5 mL of 4% Formaldehyde for 11 minutes and washed with 1×DPBS. After imaging with a Keyence BZ-X810 microscope using a 20×/0.75 objective lens, the number of cells and beads phagocytosed by each cell was counted.

#### Calcium staining assay

A 1 mM stock solution of the Fluo4 AM dye (1892-500ug, Lumiprobe) was prepared by diluting the dye in 485 µL of DMSO. To prepare a working solution, 5 µL of the stock solution was added to 995 µL of DMEM +/+ to obtain a 1 mL dye dilution per well. After 24 h of incubation, 1 mL of dye dilution was added to the wells. The Cytosoft plates were incubated at 37° C with 95% humidity and 5% CO₂ for 45 minutes. The wells were then imaged using the Keyence BZ-X810 10x/0.45 CH2 Fluor GFP at an exposure level of 1/350s.

### Statistical Analysis

All data are presented as mean ± SEM, with significance defined as P < 0.05. Analyses were performed in GraphPad Prism 6 (GraphPad Software) and in R (version 4.3.1; dplyr, ggplot2, uwot, cluster, openxlsx, stringr, patchwork, RColorBrewer, tidyr). Von Frey withdrawal thresholds were compared by Welch’s ANOVA with Dunnett’s T3 post hoc tests, and elevated zero maze, open field, and Y-maze measures by two-tailed unpaired t tests with Welch’s correction; variance-robust tests were used throughout the behavioral analyses because the assumption of equal variances was not met. For MALDI imaging, total-ion-current–normalized intensities within the hippocampus were compared between groups at the spectrum level (n = 1 animal per condition). Glial cell density and soma area were analyzed by two-way ANOVA (group × time) with Holm–Sidak post hoc tests, and the 27 single-cell morphological features were related by Pearson correlation. Cluster proportions were calculated per animal and compared between control and injured groups at each time point by Welch’s t test, with Benjamini–Hochberg correction applied across the four clusters within each time point; effect sizes are reported as Cohen’s d. Because the four cluster proportions are compositionally constrained and therefore not independent, overall cluster composition was additionally compared at each time point by permutational multivariate analysis of variance on centred log-ratio–transformed proportions, using exhaustive permutation of group labels. For the in vitro experiments, BV2 viability, morphology, engulfment, and calcium signals were analyzed by two-way ANOVA (substrate rigidity × LPS, with Sidak’s post hoc test for calcium). Exact P values and summary statistics were exported for reproducibility.

## Results

### Painful peripheral injury is paralleled by signs of mechanical sensitivity and anxiety-like behaviors

We employed a tibial fracture model of peripheral injury in mice and defined three post-surgical stages: 3 weeks (early), 7 weeks (chronic), and 20 weeks (recovery). Mechanical sensitivity was assessed using von Frey testing (Fig. 1a). Because withdrawal thresholds showed unequal variances across conditions (Brown–Forsythe test, P < 0.001 at 3 and 7 weeks), group differences were evaluated by Welch’s ANOVA with Dunnett’s T3 post hoc tests. Thresholds differed among the four paw × treatment conditions at 3 weeks (W(3, 17.9) = 26.87, p < 0.0001) and 7 weeks (W(3, 17.8) = 20.68, p < 0.0001) but not at 20 weeks (W(3, 12.7) = 2.03, p = 0.16). Ipsilateral withdrawal thresholds were markedly lower in injured than in control mice at 3 weeks (injured 0.04 g vs. control 0.65 g; mean difference 0.61 g, 95% CI [0.27, 0.96], p = 0.0010) and 7 weeks (0.03 g vs. 0.68 g; mean difference 0.65 g, 95% CI [0.22, 1.09], p = 0.0036), and no longer differed at 20 weeks (0.76 g vs. 0.89 g, p = 0.92). Contralateral thresholds did not differ between groups at any time point (all p > 0.9), indicating that the mechanical hypersensitivity was specific to the injured limb and extended beyond the period of fracture healing.

**Figure 1.**
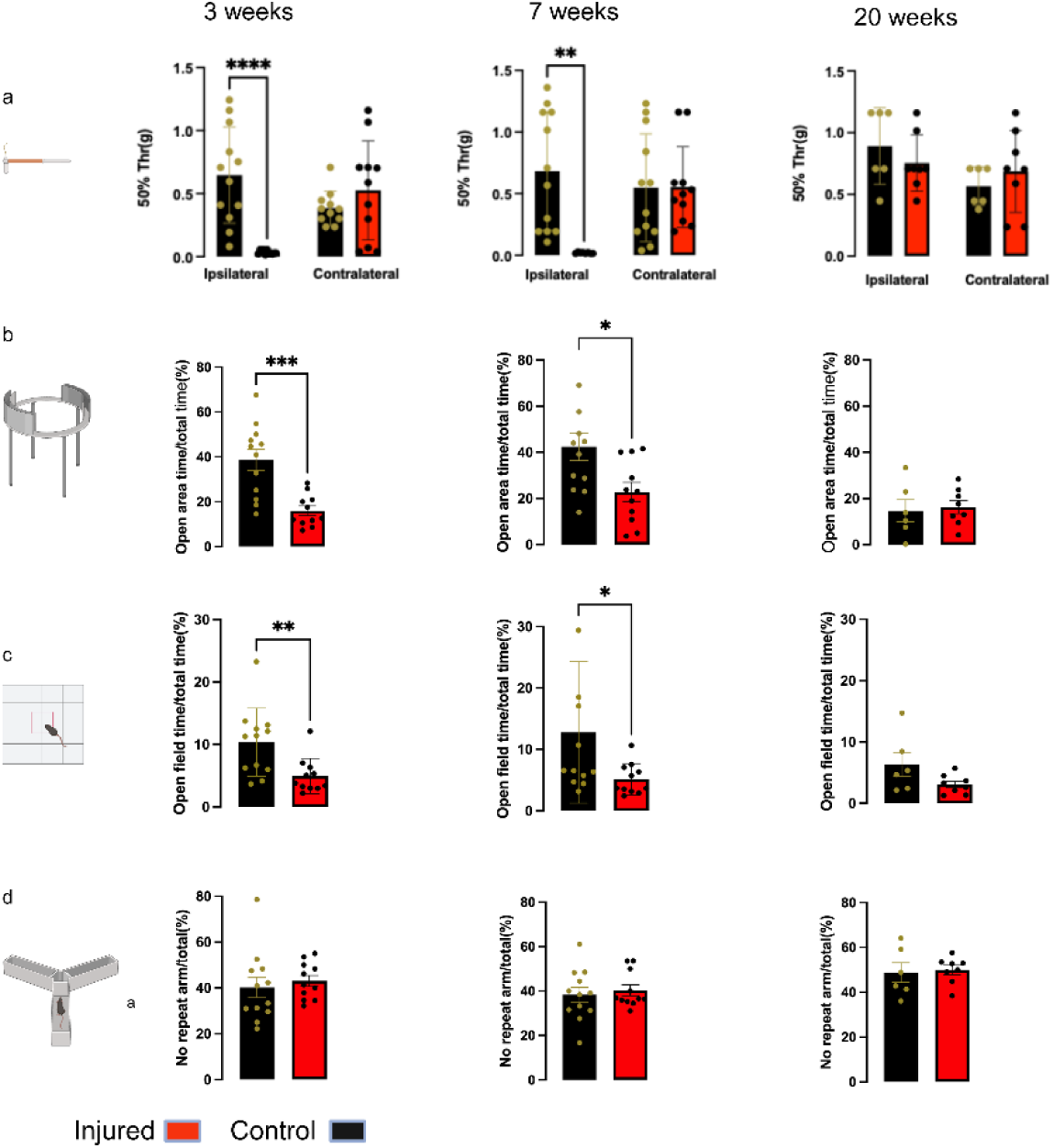
Behavioral characterization of the tibial fracture–induced chronic pain model. (a) Mechanical withdrawal thresholds (von Frey) at 3, 7, and 20 weeks post-surgery, for ipsilateral and contralateral hind paws of control and injured mice. (b) Elevated zero maze, time in the open arms (% of total). (c) Open field, time in the center zone (% of total). (d) Y-maze spontaneous alternation (% non-repeated arm entries). Von Frey thresholds were compared by Welch’s ANOVA with Dunnett’s T3 post hoc tests; zero maze, open field, and Y-maze measures by two-tailed unpaired t tests with Welch’s correction. Data are mean ± SEM and error bars indicate S.E.M.; individual points overlaid. *P < 0.05, **P < 0.01, ***P < 0.001. n = 12 (control) and 11 (injured) at 3 and 7 weeks; n = 6 (control) and 8 (injured) at 20 weeks.

Behavioral assays further revealed affective alterations during the early and chronic stages. In the elevated zero maze (Fig. 1b), injured mice spent significantly less time in the open area than controls at 3 weeks (mean difference = −22.6, 95% CI [−33.6, −11.7], t(15.3) = 4.40, p < 0.001) and 7 weeks (mean difference = −19.6, 95% CI [−34.9, −4.3], t(19.2) = 2.67, p = 0.015), with no difference at 20 weeks (mean difference = +1.5, 95% CI [−11.2, 14.3], t(8.3) = 0.28, p = 0.79). A comparable pattern was observed in the open field (Fig. 1c): injured mice spent less time in the center zone at 3 weeks (mean difference = −5.4, 95% CI [−9.2, −1.6], t(16.8) = 3.02, p = 0.008) and 7 weeks (mean difference = −7.6, 95% CI [−15.0, −0.2], t(12.2) = 2.23, p = 0.046), with no difference at 20 weeks (mean difference = −3.2, 95% CI [−8.2, 1.7], t(5.8) = 1.62, p = 0.16). In contrast, spontaneous alternation in the Y-maze (Fig. 1d) did not differ between groups at any time point (3 weeks, t(16.8) = 0.56, p = 0.58; 7 weeks, t(19.7) = 0.47, p = 0.64; 20 weeks, t(7.2) = 0.23, p = 0.82), indicating that working memory was unaffected under these conditions. Importantly, these affective differences were not accompanied by changes in general locomotor activity: neither total travel distance nor resting time differed between injured and control mice in the open field or the Y-maze at any time point (all p > 0.05; Fig. S1), indicating that the reduced exploration of anxiogenic zones in injured mice reflects an affective rather than a general locomotor phenotype. All comparisons used unpaired t tests with Welch’s correction; n = 12 (control) and 11 (injured) at 3 and 7 weeks, and n = 6 (control) and 8 (injured) at 20 weeks.

### Painful peripheral injury is paralleled by ECM abnormalities

To investigate injury-induced alterations in cellular and extracellular matrix (ECM) components within the hippocampus, we employed MALDI imaging using two complementary acquisition strategies (Fig. 2a). Intact brain sections were analyzed to profile cellular metabolites, whereas parallel sections were decellularized and subjected to tryptic digestion to enhance detection of ECM-derived peptides. Comparative analysis revealed differences in ECM-associated spectral features between injured and control mice, with injured animals exhibiting an increased number of distinct ECM-related signals (Fig. 2d). In contrast, the majority of cellular metabolite signals overlapped between groups (Fig. 2b), suggesting that detectable changes were more prominent in the ECM compartment following peripheral injury. Annotation identified 10 cellular metabolites (Fig. 2c) and four ECM proteins (Fig. 2e). Quantitative comparison of the hippocampal spectral profiles revealed higher normalized signal for glutamine and sorbitol in the injured sample, whereas the four annotated ECM-associated proteins showed lower signal intensity in the injured hippocampus. Because the MALDI analysis included one animal per condition, these observations are presented as spatially resolved molecular changes rather than inferential animal-level comparisons (Fig. S2a, b).

**Figure 2:**
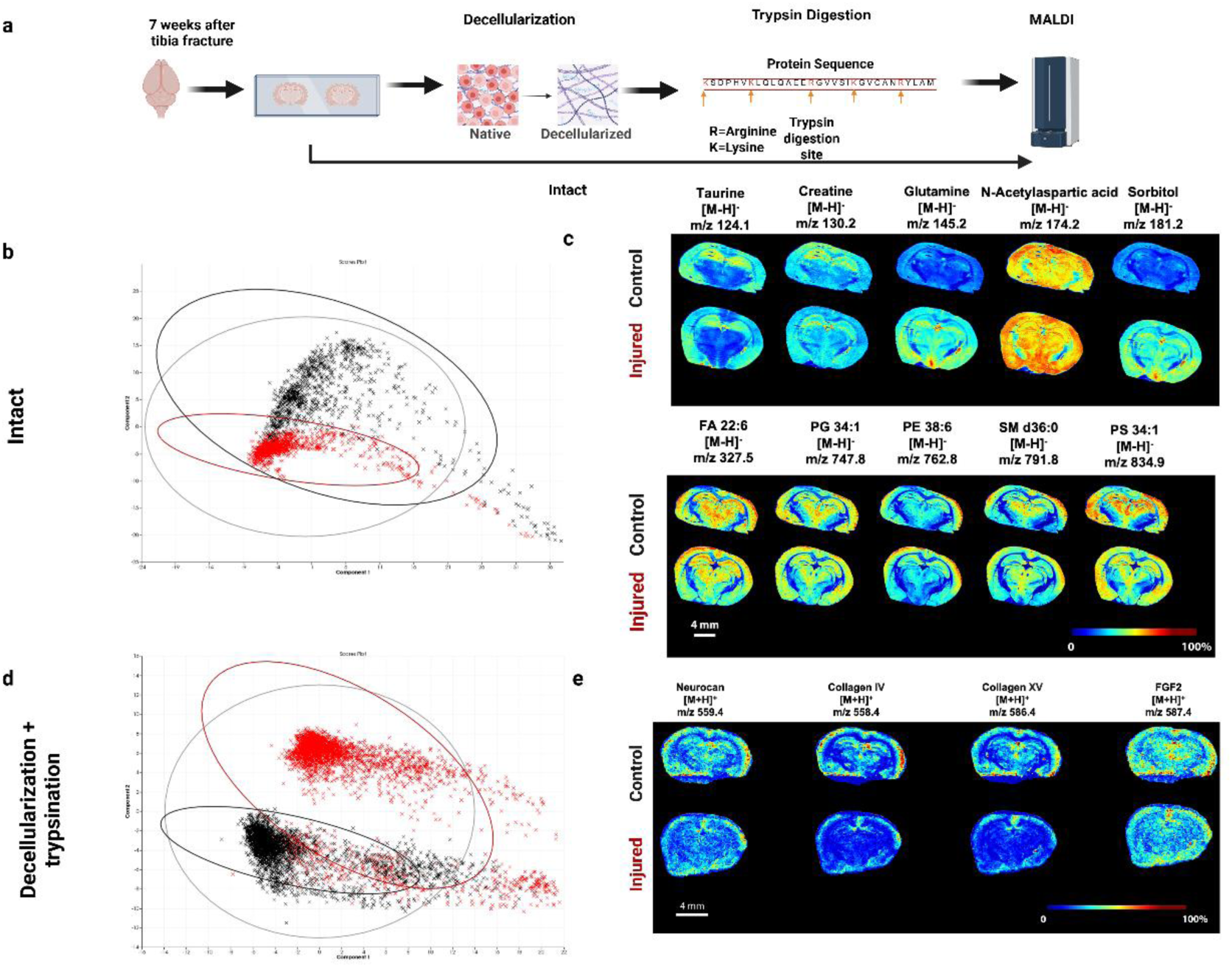
Hippocampal cellular and extracellular changes parallel to the chronic stage of peripheral injury: (a) Experimental workflow: brain slices were analyzed directly by MALDI imaging (cellular metabolites) or decellularized and trypsin-digested (ECM proteins). (b) PCA of all detected peaks from intact-brain MALDI imaging (black, control; red, injured). (c) MALDI annotation of an intact section showing metabolite distribution. (d) PCA of peaks from decellularized, trypsin-digested MALDI imaging (black, control; red, injured). (e) MALDI annotation of decellularized, trypsin-digested sections showing ECM protein distribution. Ion-intensity comparisons were made at the spectrum level. n = 1 animal per condition. Scale bars, 4 mm. MALDI, matrix-assisted laser desorption/ionization; PCA, principal component analysis; ECM, extracellular matrix; m/z, mass-to-charge ratio. Unfortunately, since n=1, we cannot include any stats here since spectra are not individual n. Please edit. You may write something like “MALDI signal distributions are shown for the hippocampal regions of one control and one injured animal. Because n = 1 animal per condition, spectral measurements are presented descriptively and were not used for animal-level inferential statistical testing.

### Peripheral injury induces temporally distinct remodeling of hippocampal microglia and astrocytes

We quantified glial density and morphology at four post-surgical time points (1, 3, 7, and 20 weeks). Analysis of microglial numbers (indicated by Iba1+ cell counts) using two-way ANOVA revealed significant main effects of time (F(3,23) = 6.92, P = 0.0017) and injury (F(1,23) = 11.93, P = 0.0022), but no significant time × injury interaction (F(3,23) = 1.22, P = 0.3238). Sidak’s multiple-comparisons test revealed a significant difference between injured and control animals at week 20, with Iba1/mouse significantly higher in injured animals (P < 0.05), whereas no significant differences were detected at week 1, 3, or 7. Astrocyte numbers (indicated by GFAP+ cell counts) were significantly higher in injured compared with control animals across the study period (main effect of injury, two-way ANOVA, F(1,23) = 12.29, P = 0.0019). There was no significant main effect of time (F(3,23) = 1.09, P = 0.3747) or injury × time interaction (F(3,23) = 0.30, P = 0.8233). Injury significantly increased both Iba1+ microglial and GFAP+ astrocytic cell numbers. For Iba1, significant main effects of injury and time were observed, whereas for GFAP only the main effect of injury was significant. Neither marker exhibited a significant time × injury interaction, indicating that the magnitude of the injury effect did not statistically differ across the examined time points. These findings suggest that microglial numbers exhibit greater temporal variability following injury, whereas the astrocytic response appears comparatively stable across the time course.

To characterize how the 27 morphological parameters relate to one another, we computed Pearson correlations across all segmented cells. Both cell types showed a modular architecture in which area-related, branching-related, and shape-related features formed internally correlated groups (Fig. S3), indicating that the 27 features vary along a small number of integrated structural dimensions. Because the features are strongly intercorrelated, we did not interpret them individually but instead used unsupervised clustering in principal component space to capture composite structural phenotypes.

To capture composite structural phenotypes, we performed unsupervised clustering in principal component space, identifying four morphological classes in microglia (Fig. 3c). Because images from the same animal are not independent, cluster proportions were compared at the level of individual animals (Welch’s t test with Benjamini–Hochberg correction; n = 4 per group, n = 3 for injured animals at 3 weeks). Microglial cluster distribution shifted in a consistent direction over time, although no individual comparison survived correction for multiple comparisons. Cluster 1 was lower in injured mice at 3 weeks (injured 45.1 ± 0.7% vs. control 55.4 ± 3.5%; d = −1.86, uncorrected p = 0.059) and 7 weeks (46.1 ± 2.1% vs. 51.6 ± 2.4%; d = −1.23, p = 0.13), and conversely higher at 20 weeks (55.6 ± 2.0% vs. 48.7 ± 2.5%; d = 1.52, p = 0.077). Clusters 3 and 4 followed the opposite trajectory: Cluster 3 was higher in injured mice at 3 weeks (16.5 ± 0.6% vs. 11.6 ± 2.1%; d = 1.49, p = 0.096) and 7 weeks (16.1 ± 1.9% vs. 12.8 ± 1.2%; d = 1.06, p = 0.19), and Cluster 4 was higher at 3 weeks (13.6 ± 1.2% vs. 9.8 ± 1.1%; d = 1.77, p = 0.069) and lower at 20 weeks (9.5 ± 1.1% vs. 13.4 ± 1.2%; d = −1.74, p = 0.049). Cluster 2 did not differ between groups at any time point, and no cluster differed at 1 week (all |d| < 0.17). Given three to four animals per group, these shifts are reported as consistent trends rather than established differences.

**Figure 3:**
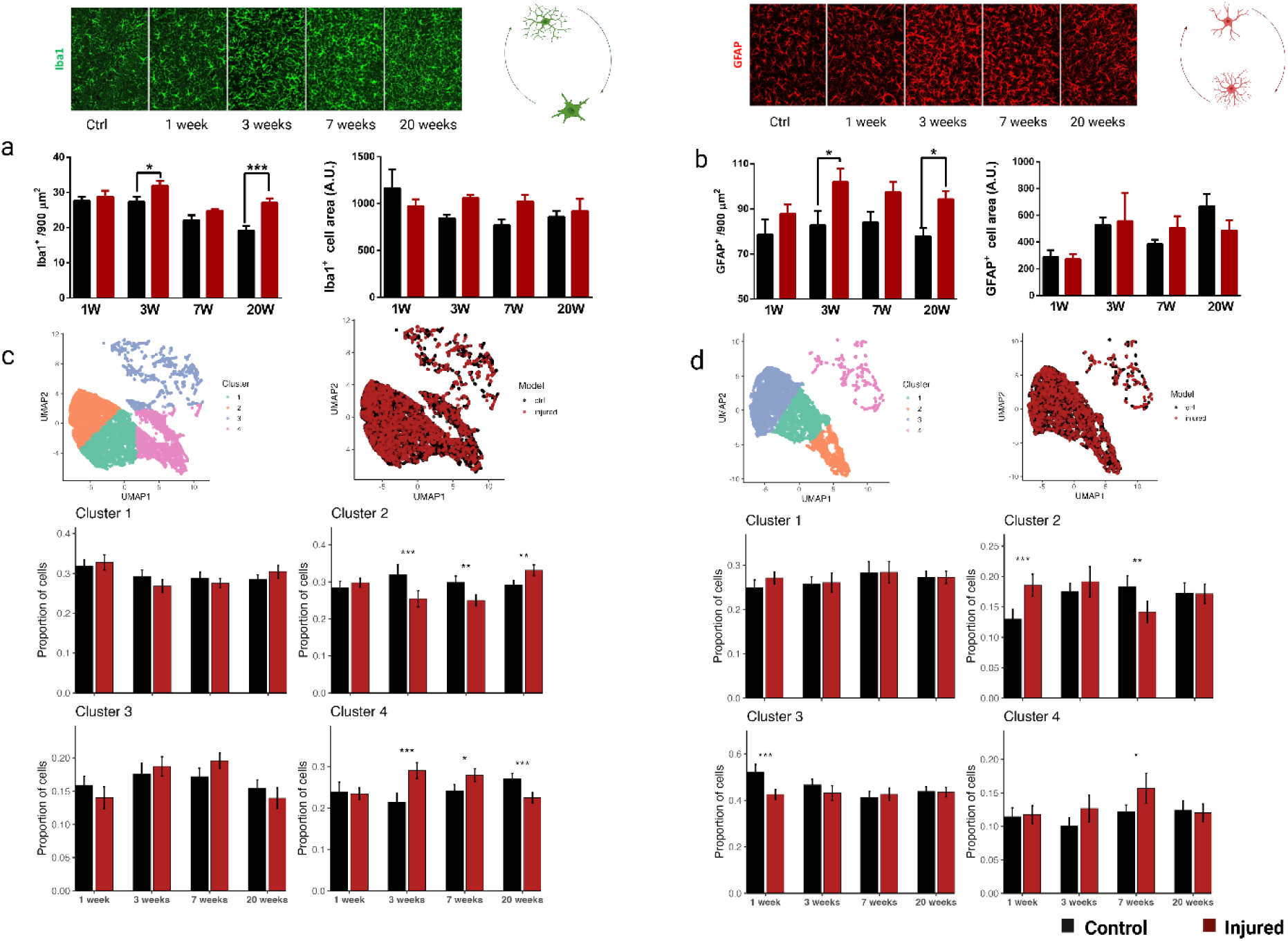
Multidimensional remodeling of glial morphology following injury: (a) Representative Iba1 immunostaining in age-matched control and at 1, 3, 7, and 20 weeks (top); quantification of Iba1⁺ cell density (cells per 900 μm²; left) and Iba1⁺ soma area (A.U.; right) (bottom). (b) GFAP immunostaining and quantification of GFAP⁺ density and soma area, as in (a). (c) UMAP of microglial features (27 per cell), colored by k-means cluster (left, k = 4) and group (right); bars show per-mouse cluster proportions at each time point (control black, injured red). (d) UMAP and cluster-proportion analysis for astrocytes, as in (c). Density and soma area were analyzed by two-way ANOVA with Holm–Sidak post hoc tests; cluster proportions by two-sided Fisher’s exact test. Data means SEM and error bars indicate S.E.M. *P < 0.05, **P < 0.01, ***P < 0.001. n = 4 mice/group at each time point. Iba1, ionized calcium-binding adapter molecule 1; GFAP, glial fibrillary acidic protein; UMAP, uniform manifold approximation and projection; A.U., arbitrary units.

Astrocytes likewise resolved into four morphological classes (Fig. 3d) but redistributed along a distinct temporal trajectory (two-sided Fisher’s exact test). Cluster 1 was reduced in injured mice at 1 week (49.2 ± 2.5% vs. control 58.0 ± 6.2%, p < 0.001), with no differences at later time points. Cluster 3 showed a biphasic pattern, being elevated in injured mice at 1 week (13.7 ± 1.4% vs. 8.8 ± 1.5%, p < 0.001) and reduced at 7 weeks (8.9 ± 0.7% vs. 12.4 ± 1.1%, p = 0.007). Clusters 2 and 4 did not differ between groups at any time point (all p > 0.05).

Together, these data show that injury was associated with shifts in glial density and redistribution across morphological clusters, with distinct temporal patterns between the two cell types—microglial cluster shifts peaking at 3–7 weeks and reversing by 20 weeks, and astrocytic shifts concentrated at the 1- and 7-week time points.

### Matrix biomechanics alters glial cell morphology and function *in vitro*

To determine whether rigidity influenced cell viability, we quantified live and dead cells using trypan blue exclusion. A chi-square test on the alive/dead contingency table revealed a statistically significant association between culture condition and viability (χ² = 35.66, df = 11, p = 0.0002). However, viability exceeded 93% under all conditions (minimum 93.9% of the cells were alive in all conditions), indicating that neither LPS nor matrix rigidity meaningfully compromised cell viability (Fig. S4).

To examine whether biomechanical features of the substrate are associated with microglial morphology, we cultured BV2 cells on type I collagen-coated substrates spanning a defined rigidity range (0.2, 2.0, 8.0, 16.0, 32.0, and 64.0 kPa), with or without LPS challenge. Two-way ANOVA revealed a significant main effect of rigidity on cell area (F(5, 12487) = 22.54, p < 0.0001), a significant main effect of LPS (F(1, 12487) = 286.2, p < 0.0001), and a significant rigidity × LPS interaction (F(5, 12487) = 13.25, p < 0.0001). The interaction reflected a differential LPS response across rigidities. LPS increased mean cell area at 0.2, 2, 8, 32, and 64 kPa, but this effect was attenuated at 16 kPa. At the population level, cells on the stiffest substrate (64 kPa) appeared more tightly clustered toward smaller areas compared to the broader distribution observed at 0.2–2 kPa (Fig. 4a). In contrast, cell roundness was unaffected by rigidity (F(5, 12487) = 0.0000245, p > 0.9999), LPS (F(1, 12487) = 0.0006, p = 0.9804), or their interaction (F(5, 12487) = 0.02455, p > 0.9999). Together, these findings indicate that microglial area is sensitive to both ECM stiffness and inflammatory stimuli, whereas microglial roundness remains stable under these conditions.

**Figure 4:**
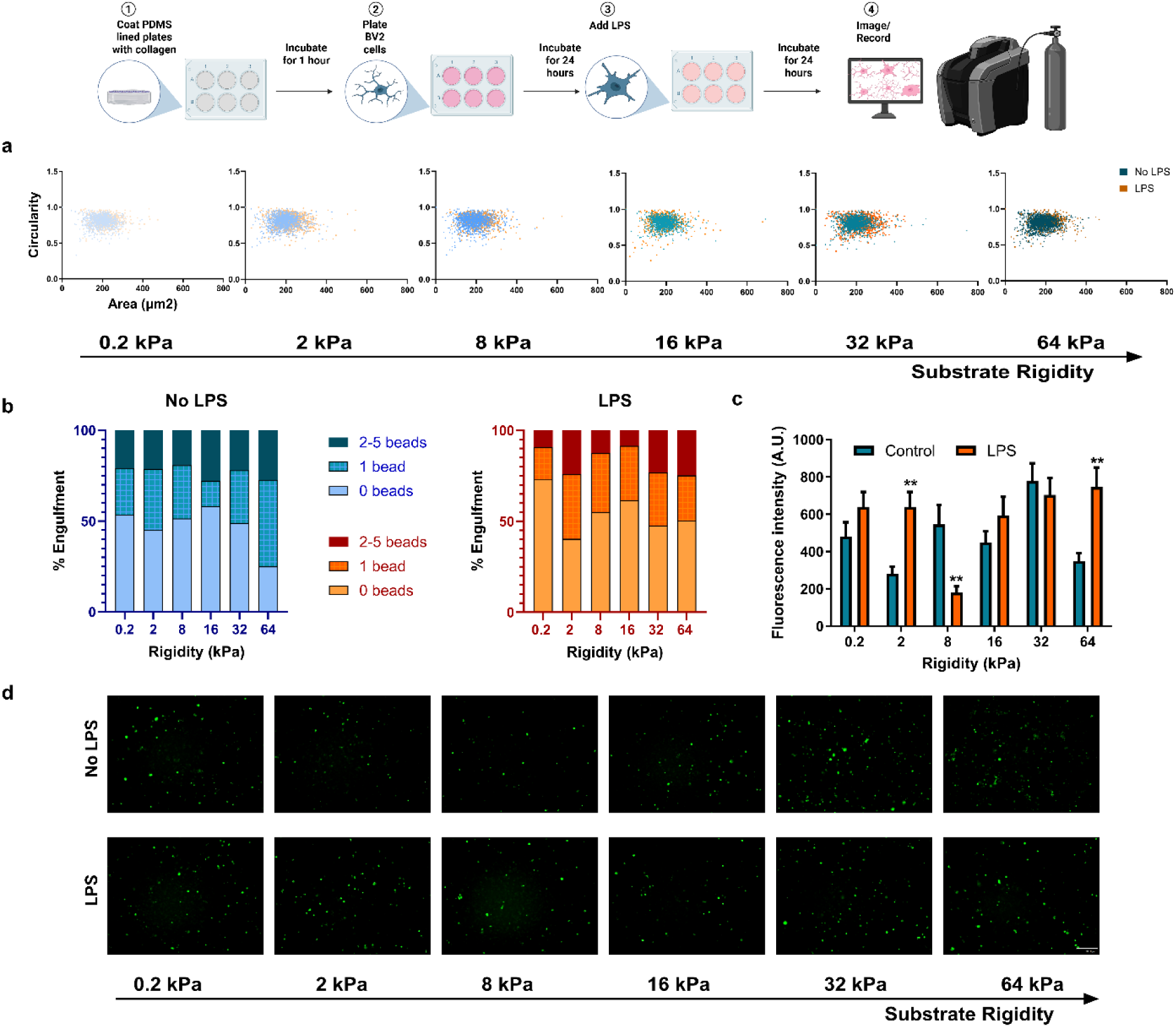
Matrix rigidity influences microglial biomechanical and functional responses. (a) Morphology: BV2 cells cultured on polydimethylsiloxane substrates (0.2–64 kPa) without LPS (teal) or with LPS (orange, 2 μg/mL, 24 h) exhibit rigidity-dependent changes in area (x-axis, soma) and circularity (y-axis, 1 = perfect circle), suggesting bidirectional modulation of microglial morphology by LPS and matrix rigidity. Data are from n = 2 biological replicates (2 technical replicates each), analyzed by two-way ANOVA. (b) Phagocytosis: Engulfment patterns of BV2 microglia cultured on PDMS substrates (0.2–64 kPa) with or without LPS (n=2) were quantified per cell and categorized as 0, 1, or 2–5 beads. Descriptive trends suggest that LPS and substrate stiffness may influence engulfment behavior, though two-way ANOVA (run separately for each engulfment category) detected no statistically significant effects (all p > 0.05). (c) Calcium signaling: Matrix rigidity and LPS were associated with changes in calcium influx in microglial cells, as measured by Fluo-4 AM imaging (n=1).

Substrate rigidity was also examined for its association with phagocytic behavior. Each cell and the number of beads it engulfed were counted and categorized as 0, 1, or 2–5 beads. Two-way ANOVA revealed no statistically significant effects of LPS treatment, substrate rigidity, or their interaction on the distribution of engulfment categories (0 beads: LPS p = 0.2566, rigidity p = 0.2429, interaction p = 0.7090; 1 bead: LPS p = 0.7417, rigidity p = 0.3408, interaction p = 0.3010; 2–5 beads: LPS p = 0.3011, rigidity p = 0.8428, interaction p = 0.8624). These preliminary observations suggest a linear relationship between substrate rigidity and bead engulfment in BV2 cells, though these trends did not reach statistical significance (Figure 4b).

**Figure 5.**
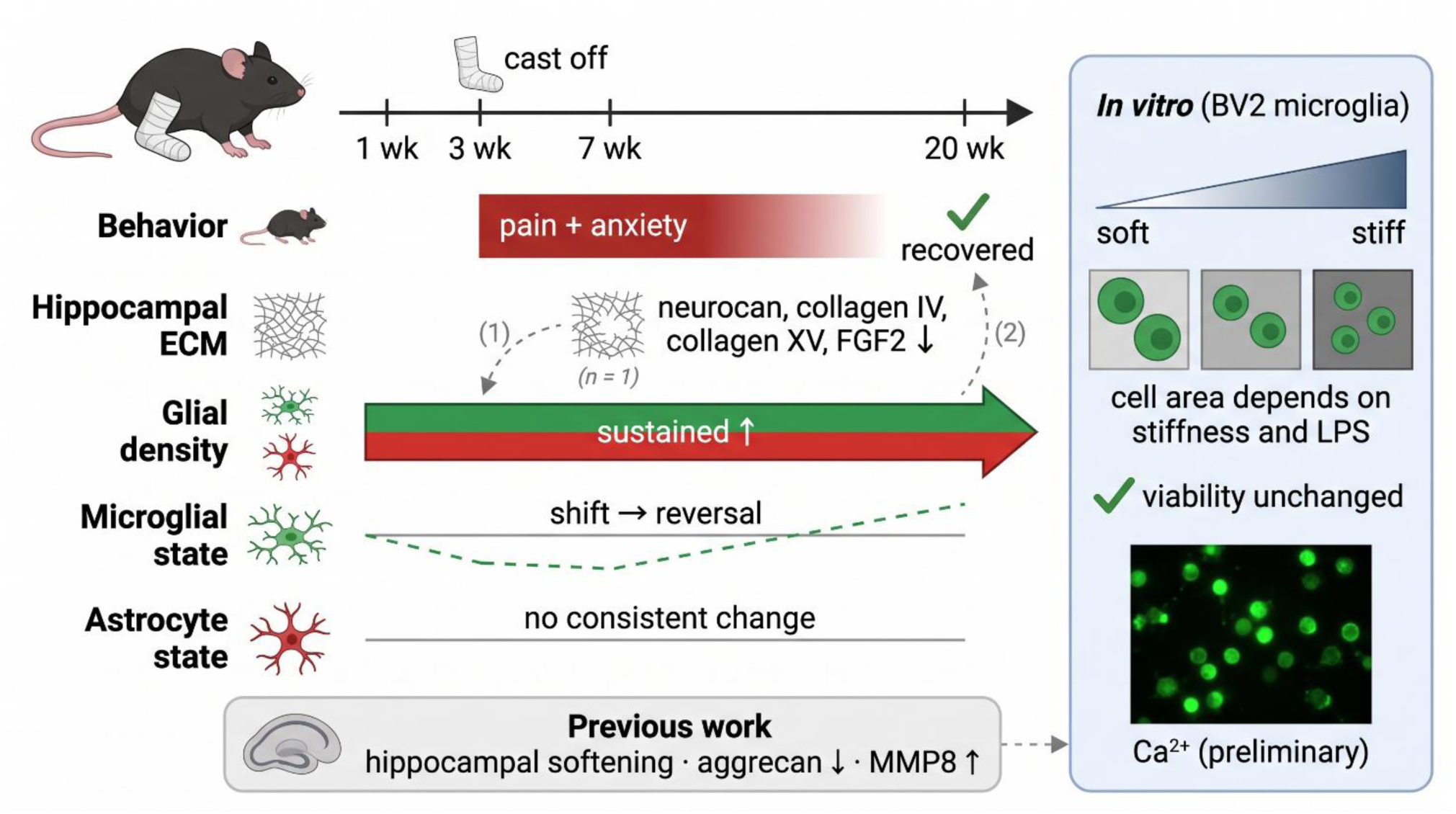
Summary of findings. Schematic overview of the behavioral, hippocampal ECM, and glial changes following tibial fracture, and of the in vitro findings in BV2 microglia. Solid elements denote observed changes; the dashed line denotes a non-significant trend. Numbered dashed arrows indicate hypothesized relationships not tested in the present study: (1) ECM changes influencing glial state and (2) glial changes contributing to the behavioral phenotype. The grey box summarizes findings previously reported in the same model [32]. The fluorescence image is schematic and does not represent experimental data. ECM, extracellular matrix; FGF2, fibroblast growth factor 2; LPS, lipopolysaccharide; MMP8, matrix metalloproteinase 8.

Calcium imaging using Fluo-4 AM was performed across the rigidity range with and without LPS. Both substrate rigidity and LPS were associated with changes in calcium signal, suggesting that mechanical input may interact with second-messenger pathways involved in microglial activation (Fig. 4d). Images were balanced for background adjustment. A two-way ANOVA with Sidak’s post hoc tests revealed significant interactions between substrate rigidity and LPS treatment in calcium signaling (F(5,3612) = 7.07, p < 0.0001). LPS treatment increased calcium activity on softer (2 kPa: mean difference = −358.9, 95% CI [-646.3, −71.55], p = 0.0061) and stiffer substrates (64 kPa: −396.4, [-639.9, −152.9], p = 0.0001), while causing the opposite effect at intermediate rigidity (8 kPa: 365.6, [64.45, 666.7], p = 0.0084). No significant differences were observed at 0.2 kPa (p = 0.5139), 16 kPa (p = 0.8142), or 32 kPa (p = 0.9810), indicating rigidity-dependent thresholds for inflammatory sensitivity. No LPS samples showed the highest baseline activity at 32 kPa (marginal mean = 777.5), whereas LPS-treated intensity peaks occurred at 8 kPa (marginal mean = 748.3). These findings indicate that calcium signaling in BV2 cells is modulated by matrix stiffness in an LPS-dependent manner, with LPS amplifying effects at both extremes of the rigidity spectrum but suppressing activity at mid-range stiffnesses (Figure 4c). Representative Fluo-4 AM images illustrate the LPS- and rigidity-dependent differences in calcium signals described above (Fig. 4d).

Together, these *in vitro* observations suggest that BV2 microglia respond to substrate rigidity across multiple functional dimensions. Morphology and calcium signaling exhibited apparent rigidity-dependent changes modulated by LPS, while phagocytosis showed descriptive trends that did not reach statistical significance. Viability was high (>93%) across all conditions and was not meaningfully affected by rigidity or LPS.

## Discussion

In this study, we examined whether chronic peripheral injury is accompanied by time-dependent changes in the hippocampal extracellular matrix and by parallel changes in glial morphology and affective behavior. Combining longitudinal behavior, MALDI-TOF imaging of intact and decellularized hippocampi at the chronic pain stage, high-dimensional glial morphometry across four time points (1, 3, 7, and 20 weeks), and an *in vitro* biomechanical assay, we observed that injury was associated with selective changes in detected hippocampal ECM protein signals at 7 weeks, with time-dependent shifts in microglial and astrocytic density and morphological cluster distribution across these time points, and with reduced exploration in two anxiety paradigms during the early and chronic stages. *In vitro*, varying substrate rigidity was associated with changes in BV2 microglial morphology and calcium signaling. Together, these results are consistent with a contribution of the hippocampal ECM-glia axis to pain-related affective comorbidity, although the present data do not by themselves establish causal relationships between ECM remodeling, glial change, and behavior.

The decellularization-coupled MALDI workflow revealed reduced signals for four ECM-associated proteins: neurocan, collagen IV, collagen XV, and FGF2 in the injured hippocampus, together with elevated glutamine and sorbitol. Neurocan is a brain-ECM chondroitin-sulfate proteoglycan whose gene (NCAN) is a shared genetic risk factor for mood and psychotic disorders, with expression concentrated in cortical and hippocampal regions[40]; its reduction echoes the decreased aggrecan and matrix rigidity we previously reported in this model[32]. Collagen IV and collagen XV are core constituents of the perivascular and basement-membrane matrix that maintains blood–brain-barrier integrity[41], so their coordinated loss may reflect remodeling of the vascular ECM compartment. FGF2 is especially pertinent: it is a heparan-sulfate-bound, matrix-sequestered growth factor that supports hippocampal neurogenesis and acts as an endogenous anxiolytic/antidepressant signal, with reduced hippocampal FGF2 linked to anxiety and mood disorders[42], raising the possibility that altered matrix-associated FGF2 may contribute to the affective phenotype. The concurrent decline of a structural proteoglycan, two basement-membrane collagens, and a matrix-tethered trophic factor is consistent with net catabolic ECM remodeling during chronification, in line with our earlier finding of elevated MMP8[32] and with the increased astrocyte density reported here, since reactive astrocytes drive both synthesis and turnover of matrix[43]. The accompanying rise in glutamine fits heightened astrocytic glutamate–glutamine cycling[44], whereas elevated sorbitol implicates the polyol pathway, a marker of osmotic and oxidative stress[45].

A decellularization-coupled strategy was well suited to detecting these targets. In intact tissue, abundant cellular lipids dominate the spectrum and suppress lower-abundance matrix signals; depleting the cellular compartment unmasks ECM-derived peptides that are otherwise obscured. Because ECM proteins are large, cross-linked, and poorly soluble, on-tissue tryptic digestion is required to generate detectable peptides, here annotated only when three or more peptides matched with congruent spatial distribution. Unlike bulk proteomics, MALDI imaging preserves spatial localization, resolving changes specifically within the hippocampus, and is label-free and multiplexed, avoiding dependence on antibodies unavailable for many ECM proteins. These strengths should be viewed alongside certain limitations: identities are MS-based annotations, quantification is relative, and imaging used a single chronic time point with one animal per condition, so orthogonal validation and additional time points will be needed. Neuroinflammation supports many persistent post-injury changes, including the release of pro-inflammatory cytokines, chemokines, complement proteins, proteinases, and reactive oxygen species [46] which, within the hippocampus, may contribute to the chronification of pain-related cognitive and affective changes.

Through dynamic physical, biochemical, and biomechanical crosstalk, the ECM shapes microglial morphology, phagocytic activity, and inflammatory responses [23], all of which can be disrupted in disease. Mechanosensitive ion channels such as Piezo1 have been proposed to mediate components of this biomechanical signaling [31]. Our *in vitro* observations are consistent with the view of the ECM not as a passive scaffold but as an instructive substrate that influences microglial behavior in a rigidity-dependent manner. Softer substrates, in the rigidity range of healthy brain tissue, supported a more rounded, less-engulfing microglial phenotype, while increasing rigidity was associated with progressive morphological change and—above approximately 2 kPa—a transition toward higher per-cell bead engulfment. This biphasic pattern is consistent with the hypothesis that altered matrix mechanics may bias microglia from a surveillance/homeostatic state toward a “fix-and-repair” activation state [47]. In such states, microglia have been linked to increased phagocytic engulfment of neuronal elements, with potential downstream consequences for synaptic pruning and neurodegeneration [48]. Whether the bead engulfment behavior we observed in BV2 cells corresponds to physiologically relevant phagocytic activity *in vivo* remains to be tested directly. The parallel changes in calcium signal raise the possibility that mechanical and inflammatory signals interact at the level of intracellular calcium dynamics, but this also requires direct mechanistic experiments to confirm.

From a translational perspective, the time course described here suggests that early interventions aimed at stabilizing matrix properties could be useful for the preservation of neuroprotective glial functions. Our previous work in this same tibia fracture model reported reduced hippocampal matrix rigidity, decreased aggrecan, and increased MMP8 in the chronic phase, and demonstrated that manipulation of the hippocampal ECM was sufficient to alter pain and pain-related memory [32]. The present study builds on that prior work in three respects. First, MALDI-TOF imaging at the chronic pain stage identified selective reductions in detected hippocampal ECM proteins in injured mice. Second, microglia and astrocytes showed shifts in density and high-dimensional morphological cluster distribution, with time courses that were not identical between the two cell types. Third, *in vitro*, BV2 cells responded to substrate rigidity along several functional readouts. Together with the prior causal evidence, these observations describe the cellular and molecular dynamics that accompany matrix remodeling in this model.

In addition to influencing neuronal plasticity, glial cells also shape the composition of the extracellular matrix itself. Our time-course data show increased numbers of GFAP⁺ astrocytes from 3 weeks and 20 weeks following injury. Because astrocytes are donors of ECM components and remodeling enzymes, an increased number of astrocytes might be expected to be accompanied by increased ECM production. However, the four ECM proteins detected by MALDI-TOF were reduced in injured animals, a pattern consistent with the matrix changes outlined above. One interpretation, consistent with these observations, is that the net hippocampal ECM state in this model reflects a balance between synthesis, proteolytic degradation, and turnover, and may shift toward catabolism when injury-driven inflammatory signaling is present. This would be in line with *in vitro* observations that reactive astrocytes synthesize both inhibitory and permissive proteoglycans [43], and with findings from spinal cord contusion in which aggrecan—a plasticity-inhibiting CSPG—decreased at the lesion site despite astrocytic upregulation [49]. Defining which proteolytic axes (for example, MMPs, ADAMTSs, heparanases) contribute to this catabolic shift in the chronic pain hippocampus is a direction for future work.

Our results also support the value of time-course analysis when investigating glia in the context of injury. A recent study of brain trauma in mice reported that distinct glial populations modify their morphology, abundance, and proliferation in a spatiotemporally orchestrated manner [50]. In our chronic pain model, microglia and astrocytes showed cluster-level shifts that did not follow identical time courses, with microglial density remaining elevated at 20 weeks—after mechanical sensitivity and anxiety-related measures were no longer different from controls. One descriptive interpretation is that some glial alterations persist beyond the time point at which the behavioral measures used here returned to control levels. The present study does not test whether such persistent changes are functionally meaningful or whether they could influence responses to subsequent challenges.

Our *in vitro* observations support a link between substrate biomechanics and microglial function, which carries potential clinical implications for the design of CNS implants [24] and for the development of non-invasive biomechanical brain imaging tools as candidate biomarkers of pain-related plasticity [51]. We used type I collagen-coated PDMS substrates because the variable of interest in this set of experiments was substrate rigidity itself, and collagen-coated PDMS is a widely used and reproducible platform for establishing defined stiffness gradients in CNS-relevant biomechanical studies. Type I collagen is not a major component of brain ECM, in which laminins, hyaluronan-binding proteoglycans, and tenascins are more abundant; convergent results from previous work using PDMS or polyacrylamide gels with collagen coatings to study CNS cell mechanosensitivity suggest that the rigidity-related component of the response is at least partly substrate-coating–independent, although coating-specific effects cannot be excluded with the present data.

Future studies will examine the direct influence of ECM composition and rigidity on glial function in ex vivo tissues from this injury model. Engulfment assays will be used to quantify pruning of distal neuronal dendrites by microglia, complementing the bead-engulfment assays reported here. We note that the state of glial activation may be linked not only to the rate of neuronal engulfment but also to the identity of the ingested material; for example, ramified microglia in the macaque retina rarely engulf synapses despite evidence of ingestion of other neuronal and glial elements [52], suggesting that selectivity of glial-neuronal interaction is itself a feature of the surveillant state. Direct *in vivo* perturbations, such as targeted enzymatic ECM degradation, MMP inhibition, or genetic manipulation of specific matrix constituents (for example, laminin β1 or tenascin-C), will be needed to test whether discrete molecular elements of the time course described here are individually required for the affective phenotype.

An important feature of the longitudinal dataset is the dissociation between behavioral recovery and persistent glial remodeling. By 20 weeks, mechanical sensitivity and anxiety-related behavioral measures were no longer significantly different between injured and control animals, whereas microglial density remained elevated and aspects of the morphological distribution remained altered. These findings suggest that resolution of the behavioral phenotype does not necessarily coincide with complete normalization of hippocampal glial architecture. Whether these persistent changes represent residual adaptation, compensation, or altered preparedness for subsequent challenge remains to be determined.

### Limitations

Several limitations of the present study should be noted. First, *in vitro* mechanistic experiments were performed using BV2 cells rather than primary glial cultures. BV2 is a widely used murine microglial line that shares many functional features with primary microglia, including responsiveness to inflammatory stimuli, and offers reproducibility and the throughput needed for the multi-rigidity, multi-readout experiments performed here[53, 54]. BV2 cells nonetheless differ from primary microglia in transcriptomic profile, baseline activation tone, and responsiveness to immune stimuli[55, 56]. Validation in primary microglia and, where feasible, in iPSC-derived human microglia will be useful in future work. Second, the *in vitro* biomechanical platform tested only type I collagen-coated substrates, while brain ECM is composed predominantly of laminins, hyaluronan-binding proteoglycans, and tenascins; future experiments using brain-relevant matrix coatings will help define the contribution of physiologically relevant biochemical cues. Third, the present study employed only male mice; we have previously reported sex-related differences in pain-related cognition in this model [22], and the affective and ECM-glia patterns described here may differ in female animals, motivating sex-disaggregated analysis in follow-up work. Finally, MALDI-TOF imaging was performed at a single time point (7 weeks, chronic pain stage); a more comprehensive view of ECM dynamics across the chronification trajectory would benefit from MALDI sampling at additional time points and from complementary biochemical analyses.

## Acknowledgements

MT was supported by NIH grant RNS140306.

