## Supplementary material for "Hippocampal extracellular matrix remodeling and temporally distinct glial reorganization following peripheral injury": supplymental figure

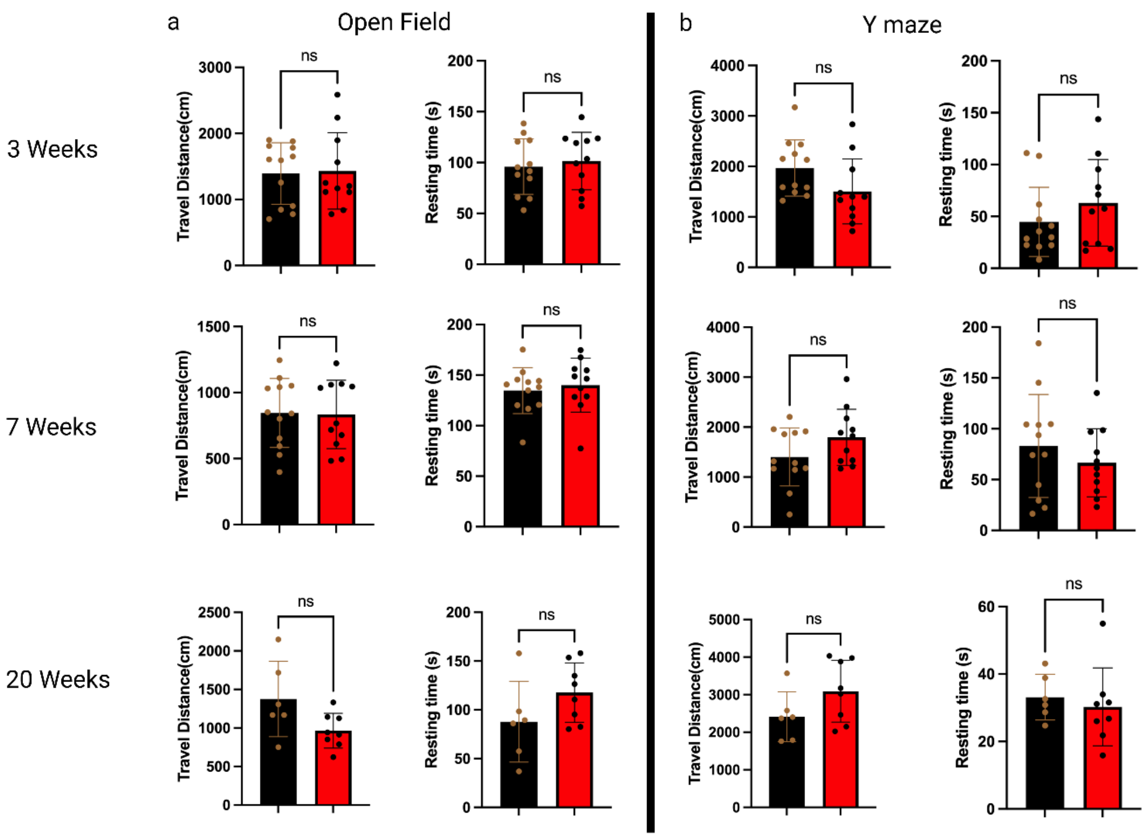


**Figure S1. Locomotor activity is unchanged after injury in the open field and Y-maze.** Total travel distance (cm) and resting time (s) for control (black) and injured (red) mice in the open field (a) and Y-maze (b) at 3, 7, and 20 weeks. Neither measure differed between groups in either apparatus at any time point. Comparisons by two-tailed unpaired t tests with Welch's correction (ns, $P \geq 0.05$). Data means SEM and error bars indicate S.E.M.; individual points overlaid. n = 12 (control) and 11 (injured) at 3 and 7 weeks; n = 6 (control) and 8 (injured) at 20 weeks. The absence of locomotor differences indicates that the reduced exploration of anxiogenic zones (Fig. 1b,c) reflects an affective rather than a locomotor phenotype.

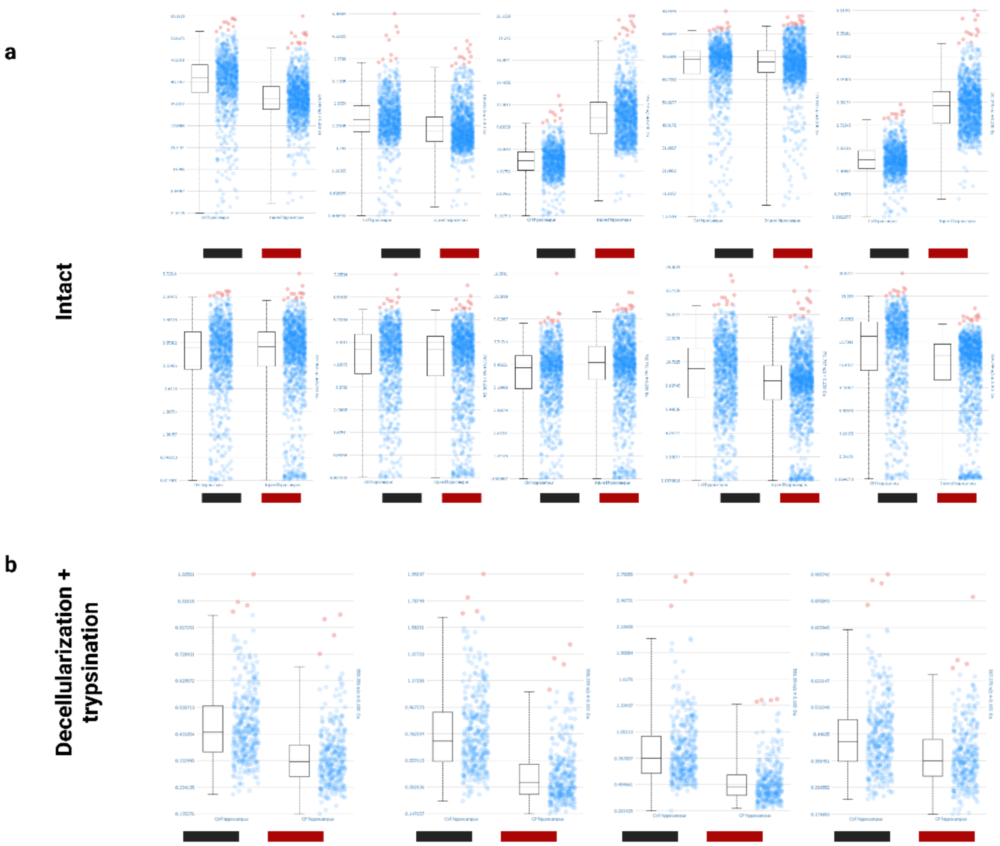


**Figure S2: Chronic Pain-Induced Changes in Mouse Hippocampal Metabolites and Extracellular Matrix Proteins.** (a) Intact MALDI annotation of hippocampal metabolites (black, control; red, injured). (b) MALDI annotation of decellularized, trypsin-digested hippocampal slices showing ECM proteins (black, control; red, injured). Box plots show median, interquartile range, and 0–99% whiskers, with individual spectra overlaid. n = 1 animal per condition. MALDI, matrix-assisted laser desorption/ionization; ECM, extracellular matrix; m/z, mass-to-charge ratio.

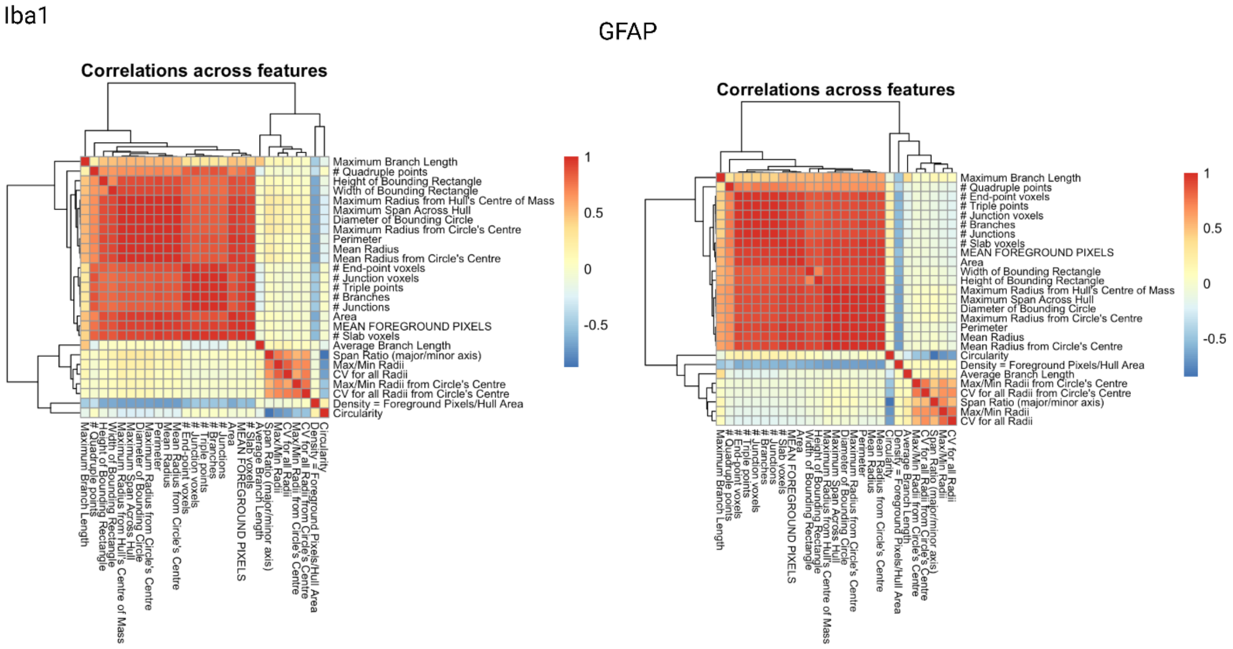


**Figure S3. Correlation matrices of 27 morphological features in microglia and astrocytes.** Pearson correlation matrices of the 27 per-cell morphological parameters, with hierarchical-clustering dendrograms on the top and left axes. (a) Microglia (Iba1$^+$). (b) Astrocytes (GFAP$^+$). Color scale: red, positive (to +1); blue, negative (to ≈ −0.5); yellow/cream, ~0. Three internally correlated groups are evident in both cell types: a size/branching block (Area, Perimeter, Number of Branches, Number of Junctions, Diameter of Bounding Circle, Mean Foreground Pixels); a radius-variability/span cluster (Span Ratio, Max/Min Radii, CV for all Radii); and shape features (Circularity, Density). Values are Pearson correlation coefficients computed across all segmented cells. Iba1, ionized calcium-binding adapter molecule 1; GFAP, glial fibrillary acidic protein; CV, coefficient of variation.

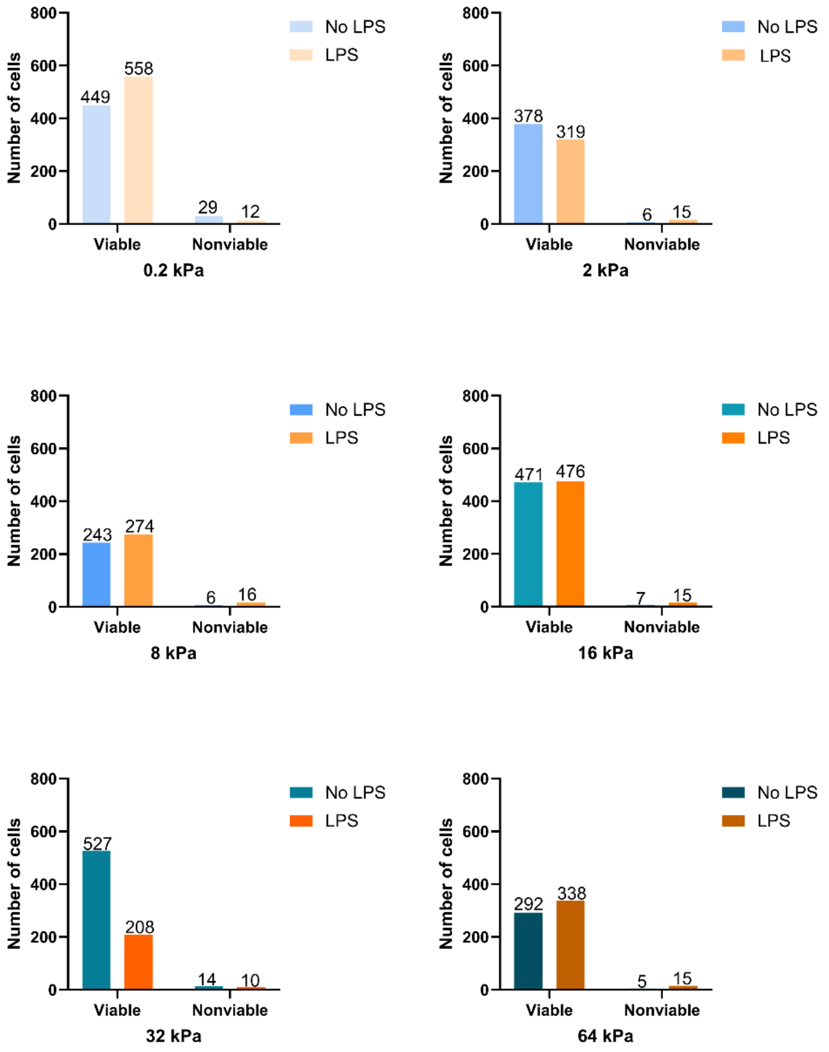


**Figure**

**Figure S4. BV2 viability across substrate rigidities and LPS treatment.** BV2 cells were cultured on type I collagen-coated PDMS substrates (0.2–64 kPa) with or without LPS (2 µg/mL, 24 h), and viability was assessed by trypan blue exclusion. Bars show the numbers of viable and nonviable cells in each condition. Viability exceeded 93% in all conditions (range 93.9–98.5%) and did not vary systematically with substrate rigidity. The proportion of nonviable cells differed across the 12 conditions ($\chi^2 = 35.66$, df = 11, p = 0.0002), but the absolute differences were small ($\leq$ 4.6 percentage points).
